## Supplementary information for "Differences in the regulation mechanisms of the glutamine synthetase from methanogenic archaea unveiled by structural investigations"

**This PDF file includes:**

Tables S1 and S2

Figures S1 to S15

SI references

**Other supporting materials for this manuscript include the following:**

Excel file of Table S3

Movie "*Mt*GS_conformational_changes.mp4"

**Table S1. X-ray analysis statistics for *Mt*GS.**

| | *Mt*GS-apo-TbXo4 | *Mt*GS-apo without TbXo4 | *Mt*GS-2OG/$Mg^{2+}$/ATP | *Mt*GS-2OG/$Mg^{2+}$ |
|---|---|---|---|---|
| **Data collection** | | | | |
| Wavelength (Å) | 0.97625 | 1.00004 | 1.30511 | 0.99999 |
| Space group | $C222_1$ | $C222_1$ | *P*1 | *P*1 |
| Resolution (Å) | 49.78 – 1.65 (1.74 – 1.65) | 76.67 – 2.43 (2.57 – 2.43) | 110.03 – 2.15 (2.30 – 2.15) | 203.54 – 2.91 (3.12 – 2.91) |
| Cell dimensions | | | | |
| a, b, c (Å) | 131.43, 228.45, 204.80 | 132.65, 230.24, 205.59 | 112.34, 131.77, 131.51 | 131.81, 131.93, 203.54 |
| α, β, γ (°) | 90, 90, 90 | 90, 90, 90 | 60.04, 87.72, 67.34 | 89.95, 89.86, 60.05 |
| $R_{merge}$(%)[a] | 6.3 (91.8) | 26.9 (179.5) | 11.0 (95.0) | 12.2 (50.1) |
| $R_{pim}$ (%)[a] | 3.9 (56.6) | 7.3 (46.4) | 6.8 (59.0) | 7.6 (34.4) |
| $CC_{1/2}$ [a] | 0.999 (0.782) | 0.998 (0.782) | 0.995 (0.569) | 0.993 (0.548) |
| I/σ*I*[a] | 11.9 (1.6) | 7.4 (1.5) | 9.3 (1.3) | 5.9 (1.6) |
| Spherical completeness[a] | 98.9 (97.9) | 86.6 (28.7) | 76.9 (21.1) | 64.8 (16.8) |
| Ellipsoidal completeness[a] | / | 93.3 (49.5) | 92.2 (58.8) | 91.7 (69.3) |
| Redundancy[a] | 6.9 (6.9) | 14.5 (15.6) | 3.6 (3.4) | 3.5 (3.0) |
| Nr. unique reflections[a] | 361,584 (51,999) | 102,291 (5,117) | 248,419 (12,420) | 169,224 (8,461) |
| **Refinement** | | | | |
| Resolution (Å) | 49.78 – 1.65 | 58.89 – 2.43 | 42.88-2.15 | 43.74 – 2.91 |
| Number of reflections | 361148 | 102,286 | 248,376 | 169,113 |
| $R_{work}/R_{free}$[b] (%) | 16.30/18.50 | 22.94/26.82 | 17.10/20.30[d] | 25.41/27.80 |
| Number of atoms | | | | |
| Protein | 21,321 | 21,082 | 42,408 | 84,792 |
| Ligands/ions | 81 | 154 | 710 | 371 |
| Solvent | 2,932 | 293 | 3,041 | 0 |
| Mean B-value (Å$^2$) | 33.52 | 52.65 | 38.36 | 63.17 |
| Molprobity clash score | 2.32 | 4.01 | 1.99 | 1.09 |
| Ramachandran plot | | | | |
| Favored regions (%) | 98.20 | 96.51 | 97.92 | 96.26 |
| Outlier regions (%) | 0 | 0.04 | 0.22 | 0.24 |
| rmsd[c] bond lengths (Å) | 0.012 | 0.004 | 0.009 | 0.010 |
| rmsd[c] bond angles (°) | 1.485 | 0.654 | 0.93 | 1.274 |
| **PDB ID code** | 8OOL | 8OON | 8OOO | 8OOQ |

[a] Values relative to the highest resolution shell are within parentheses. [b] Rfree was calculated as the Rwork for 5 % of the reflections that were not included in the refinement. [c] rmsd, root mean square deviation. [d] Rfactor and Rfree are from the PDB validation report.

**Table S2. X-ray analysis statistics for *Ms*GS.**

| | *Ms*GS apo 1 | *Ms*GS apo 2 | *Ms*GS-$Mg^{2+}$/ATP |
|---|---|---|---|
| **Data collection** | | | |
| Wavelength (Å) | 1.00004 | 1.00002 | 0.97949 |
| Space group | $P2_1$ | $P4_332$ | $P2_1$ |
| Resolution (Å) | 109.99 - 2.64 (2.93 - 2.64) | 130.75 - 3.09 (3.17 - 3.09) | 78.85 - 2.70 (2.88 - 2.70) |
| Cell dimensions | | | |
| a, b, c (Å) | 130.92, 195.65, 133.44 | 226.46, 226.46, 226.46 | 131.55, 197.40, 135.17 |
| α, β, γ (°) | 90, 94.71, 90 | 90, 90, 90 | 90, 94.89, 90 |
| $R_{merge}$(%)[a] | 7.6 (87.0) | 34.5 (309.9) | 6.2 (68.8) |
| $R_{pim}$ (%)[a] | 3.5 (46.7) | 6.8 (60.7) | 3.3 (37.0) |
| $CC_{1/2}$ [a] | 0.999 (0.600) | 0.998 (0.486) | 0.999 (0.701) |
| I/σ/[a] | 15.5 (1.6) | 12.2 (1.4) | 17.4 (2.0) |
| Spherical completeness[a] | 62.8 (11.5) | 96.6 (63.9) | 68.5 (19.8) |
| Ellipsoidal completeness[a] | 90.3 (78.7) | 96.6 (63.8) | 88.1 (89.8) |
| Redundancy[a] | 5.7 (4.4) | 26.6 (26.8) | 4.3 (4.4) |
| Nr. unique reflections[a] | 123,574 (6,180) | 35,711 (1,793) | 128,572 (6,429) |
| **Refinement** | | | |
| Resolution (Å) | 62.96 - 2.64 | 65.37 - 3.09 | 49.10 - 2.70 |
| Number of reflections | 123,529 | 35,696 | 128,533 |
| $R_{work}/R_{free}$[b] (%) | 19.14/22.52 | 18.63/21.62 | 19.61/21.72 |
| Number of atoms | | | |
| Protein | 40,821 | 6,920 | 40,815 |
| Ligands/ions | 76 | 64 | 431 |
| Solvent | 66 | 0 | 33 |
| Mean B-value ($Å^2$) | 72.24 | 80.11 | 70.78 |
| Molprobity clash score | 2.19 | 2.30 | 2.60 |
| Ramachandran plot | | | |
| Favored regions (%) | 98.14 | 96.31 | 97.95 |
| Outlier regions (%) | 0 | 0 | 0.02 |
| rmsd[c] bond lengths (Å) | 0.011 | 0.004 | 0.011 |
| rmsd[c] bond angles (°) | 1.35 | 0.678 | 1.348 |
| **PDB ID code** | 8OOW | 8OOX | 8OOZ |

[a] Values relative to the highest resolution shell are within parentheses. [b] Rfree was calculated as the Rwork for 5 % of the reflections that were not included in the refinement. [c] rmsd, root mean square deviation.

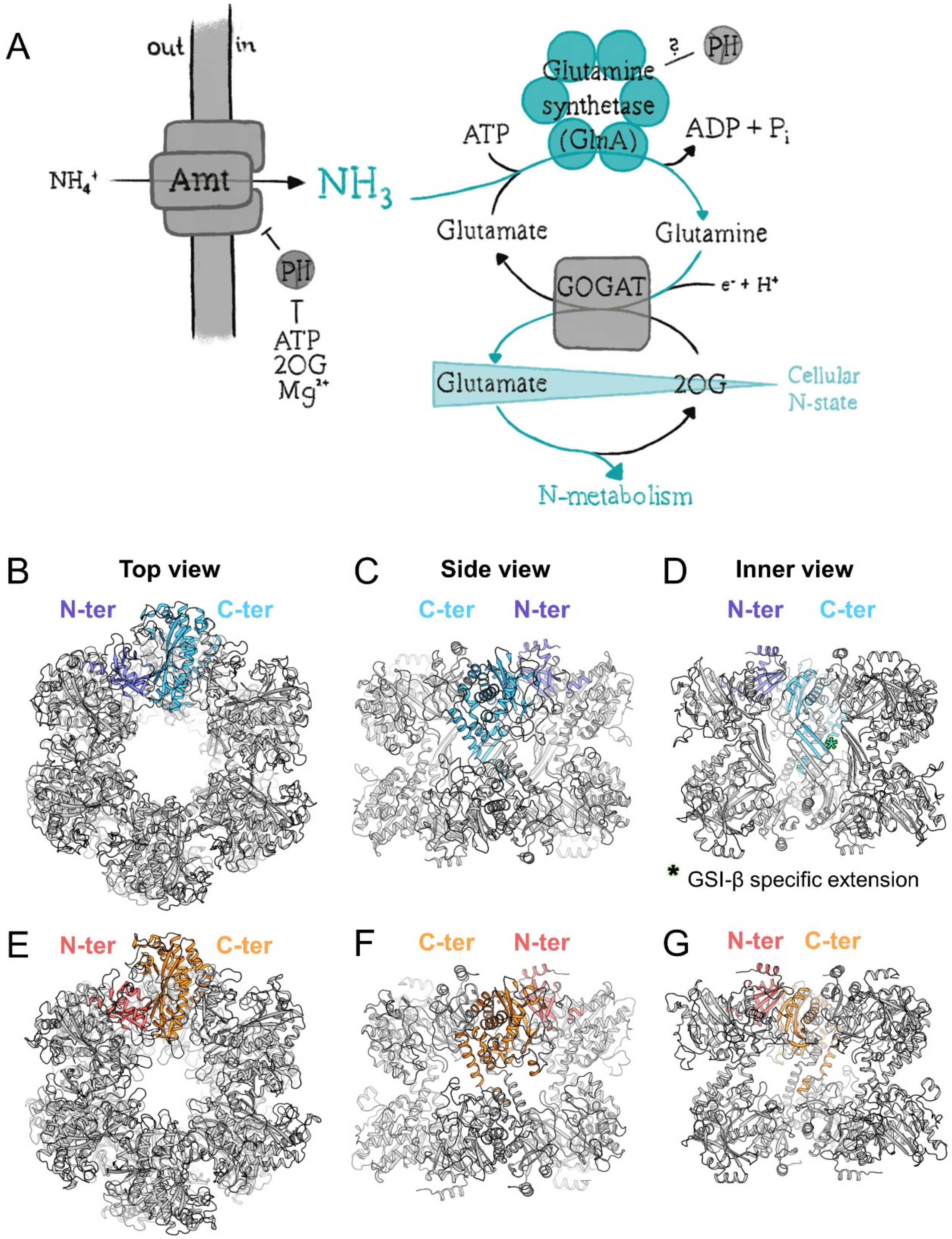


**Figure S1: Physiological role of GS-GOGAT in methanogens and structural organization of GSI-α and GSI-β.** (A) GS-GOGAT reaction scheme. 2OG concentration increases under nitrogen starvation, while it decreases when cellular nitrogen is available. (B-G) Comparison of homododecameric GSI-β from *Escherichia coli* (B-D, PDB 7W85) and GSI-α from *B. subtilis* (E-G, PDB 4LNN). The proteins are in cartoon with the highlighted N-terminal (dark blue and red) and C-terminal (light blue and orange). (B and E) Top view of the dodecameric GS. (C and F) outer-side view. (D and G) Inner-side view. An asterisk marks the extension characterizing the GSI-β.

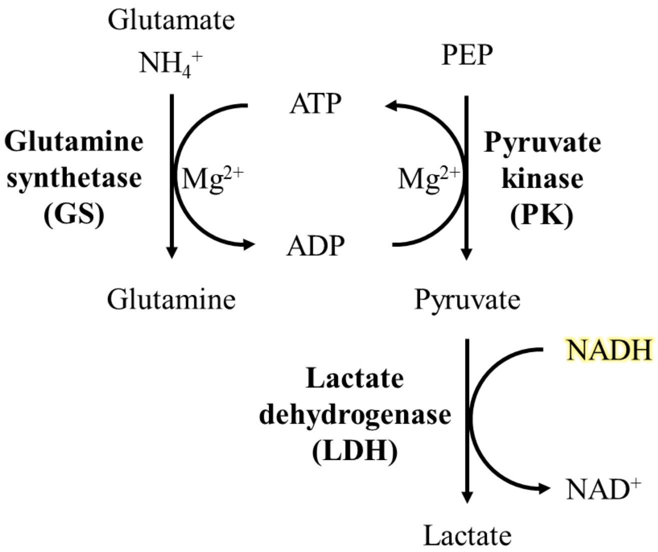


**Figure S2: Illustration of the coupled enzyme assay used in this study.** The reaction was tracked by following NADH oxidation at 340 nm. Enzyme names are bold. PEP stands for phosphoenolpyruvate.

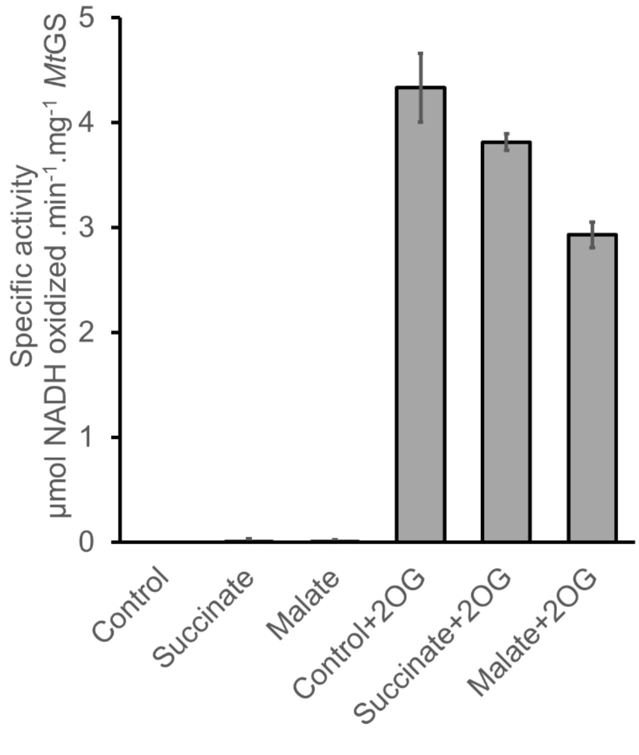


**Figure S3: 2OG specificity of *Mt*GS.** Specific activity of *Mt*GS after incubation with succinate or malate (15 mM, final first three columns) and after 20 min incubation with succinate and malate and subsequent 2OG addition (2 mM 2OG, last three columns) n=3.

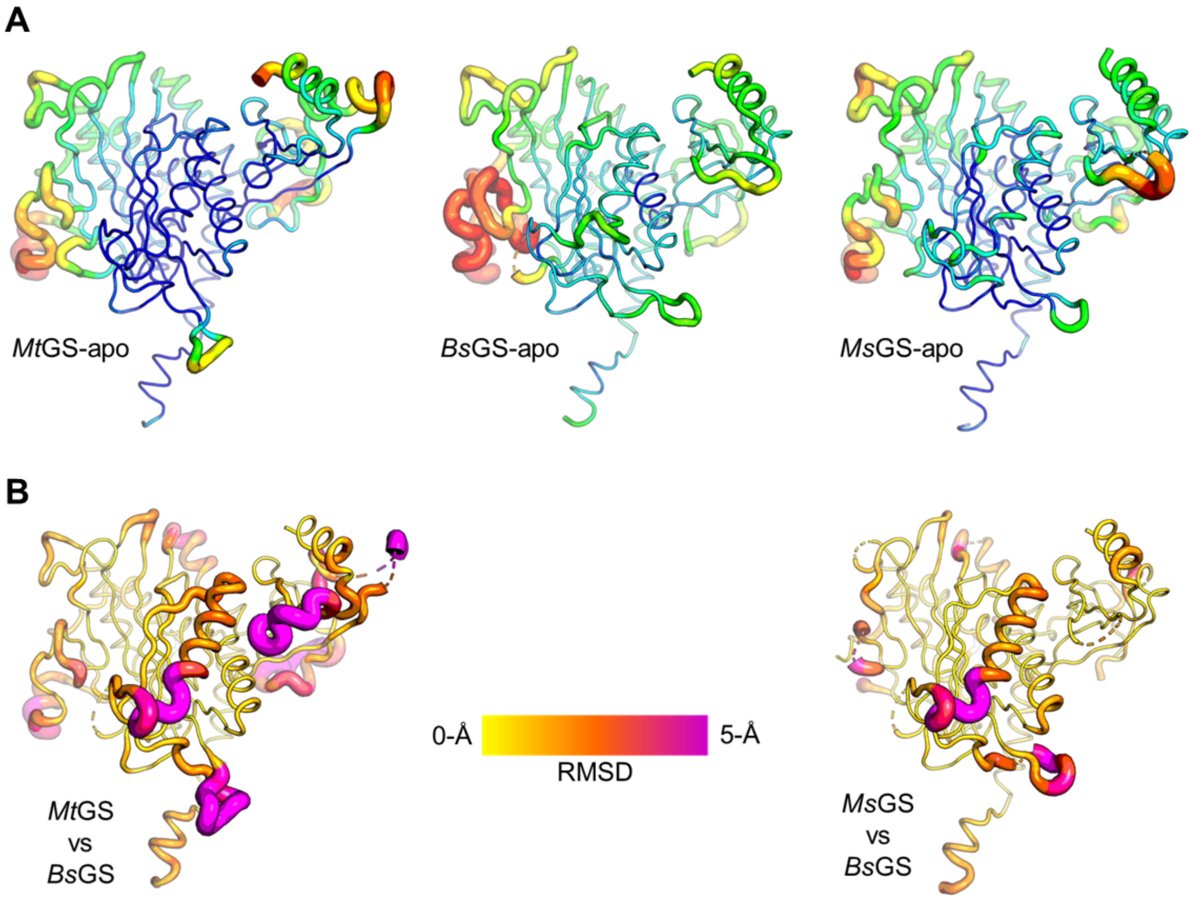


**Figure S4: Structural variation between *Ms*GS/*Mt*GS and *Bs*GS apo.** (A) B-factors and (B) RMSD values displayed as cartoon putty, for the latter the average of all chains is displayed. Average RMSD values were generated from aligning all chains of *Mt*- or *Ms*GS on chain A of *Bs*GS apo (PDB 4LNN).

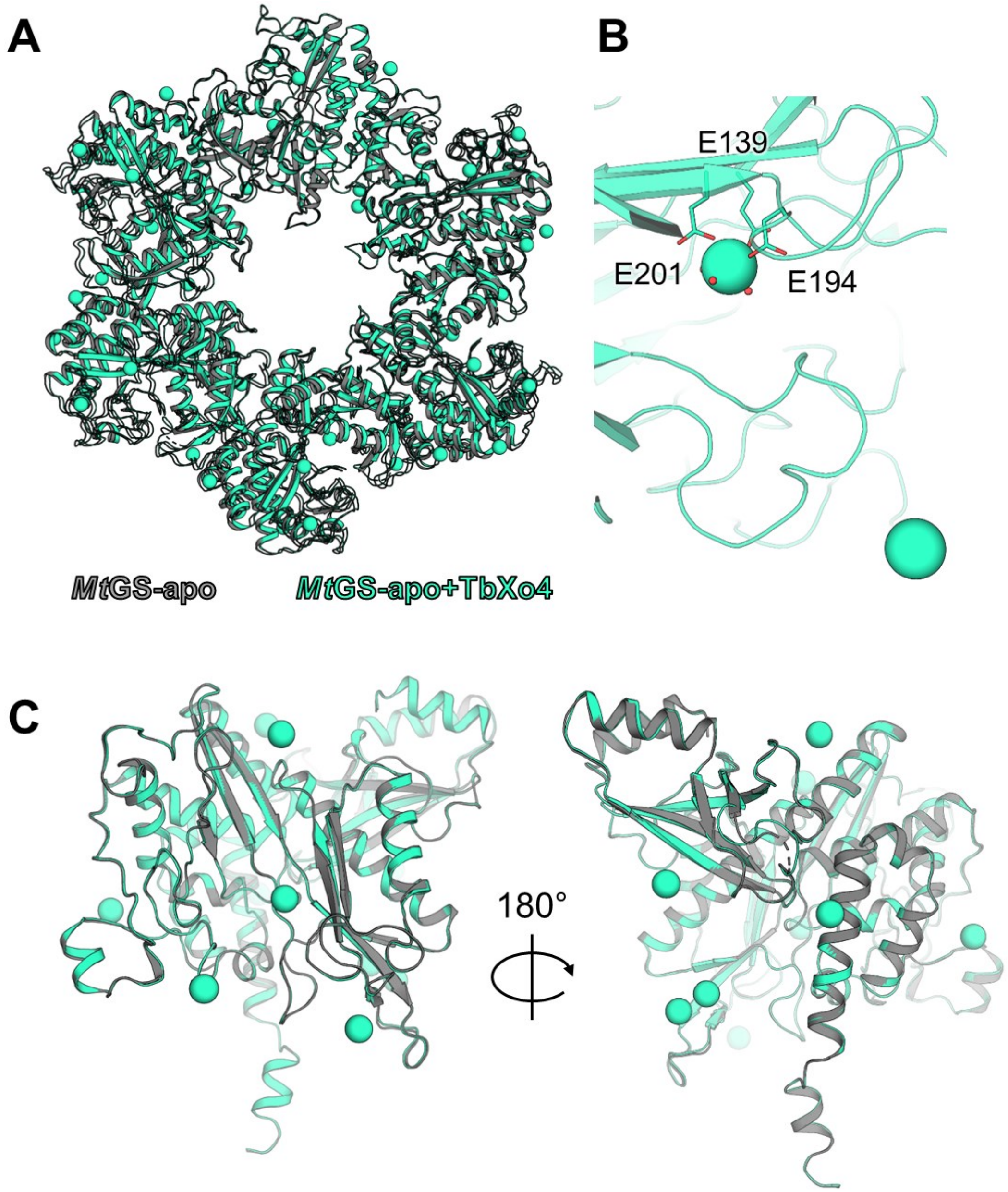


**Figure S5: Differences between *Mt*GS apo state with or without TbXo4**. (A) Overview of the hexamer from the top. (B) Binding of Tb in the active site. (C) Overview of the superposed monomers with the same pose as in Fig. 2C. For A-C, models are represented in cartoon with Tb (cyan) and $H_2O$ as spheres and the coordinating residues as sticks with oxygen atoms in red.

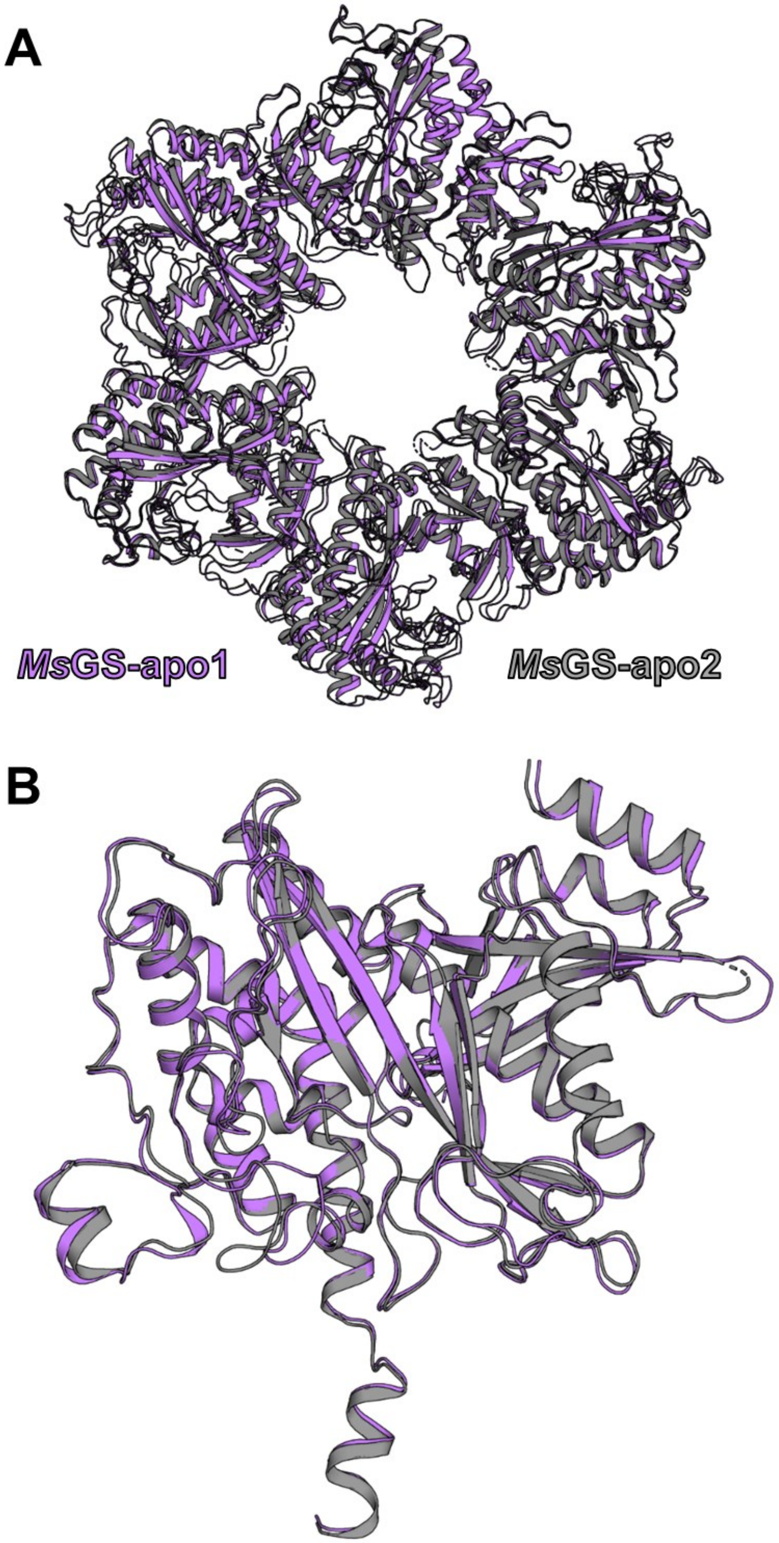


**Figure S6: Differences between the two apo forms of *Ms*GS**. (A) Superposition of the *Ms*GS apo1 and apo2 hexamers. (B) Superposition of the monomers. All models are represented as cartoons with the same pose as in Fig. 2C.

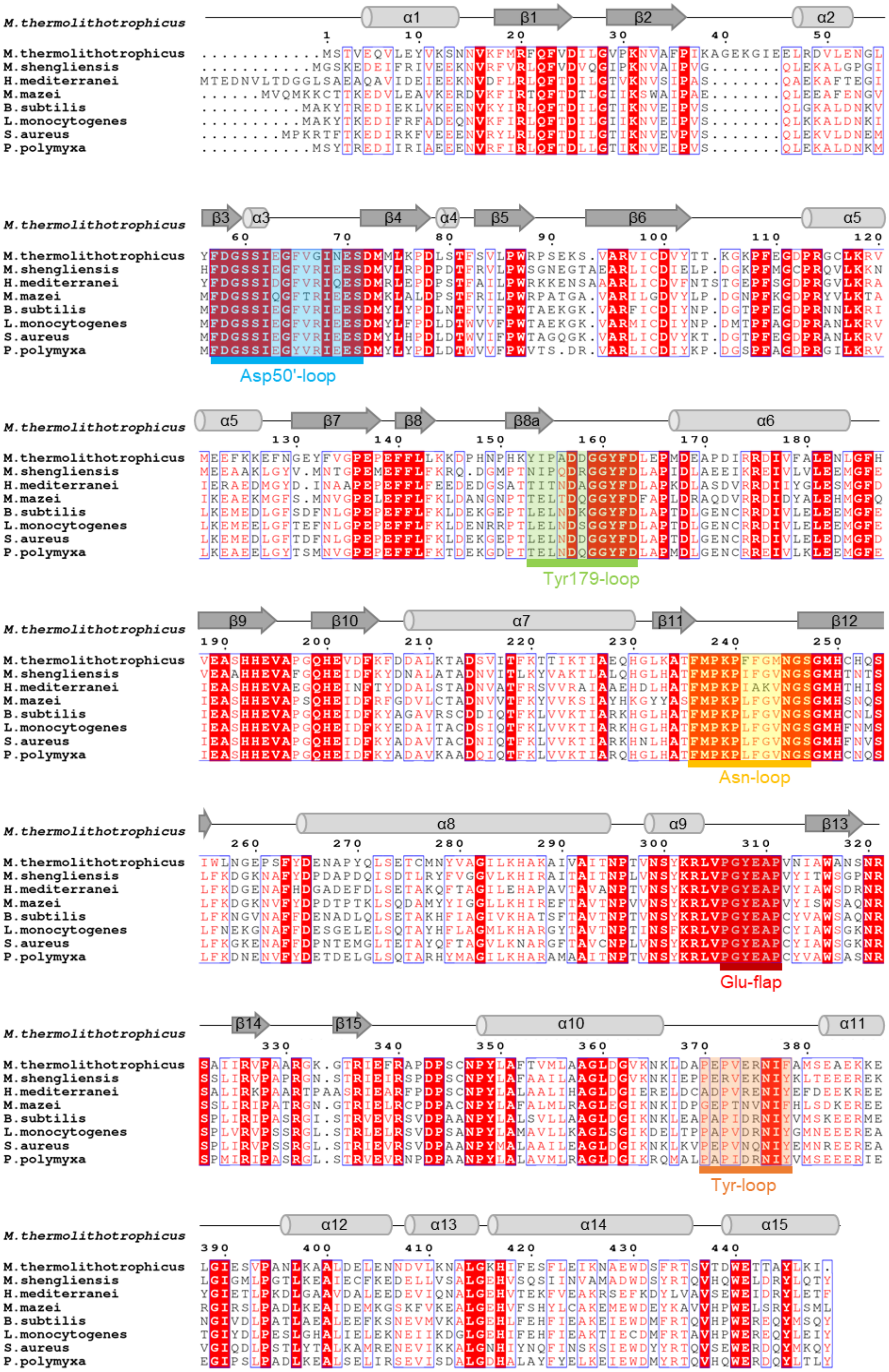

**Figure S7: Sequence conservation superposed to the secondary structure.** The secondary structure is annotated based on Travis et. al 2022 [1]. Relevant loops are labelled. WP numbers from top to bottom: WP_018154487.1, WP_042685700.1, CAR67815.1 (*H. mediterranei* GS3), WP_011032914.1 (*M. mazei* GS1), NP_389628.1, EAC9051058.1 (*Listeria monocytogenes*), WP_086038154.1 (*Staphylococcus aureus*), WP_016822091.1 (*Paenibacillus polymyxa*).

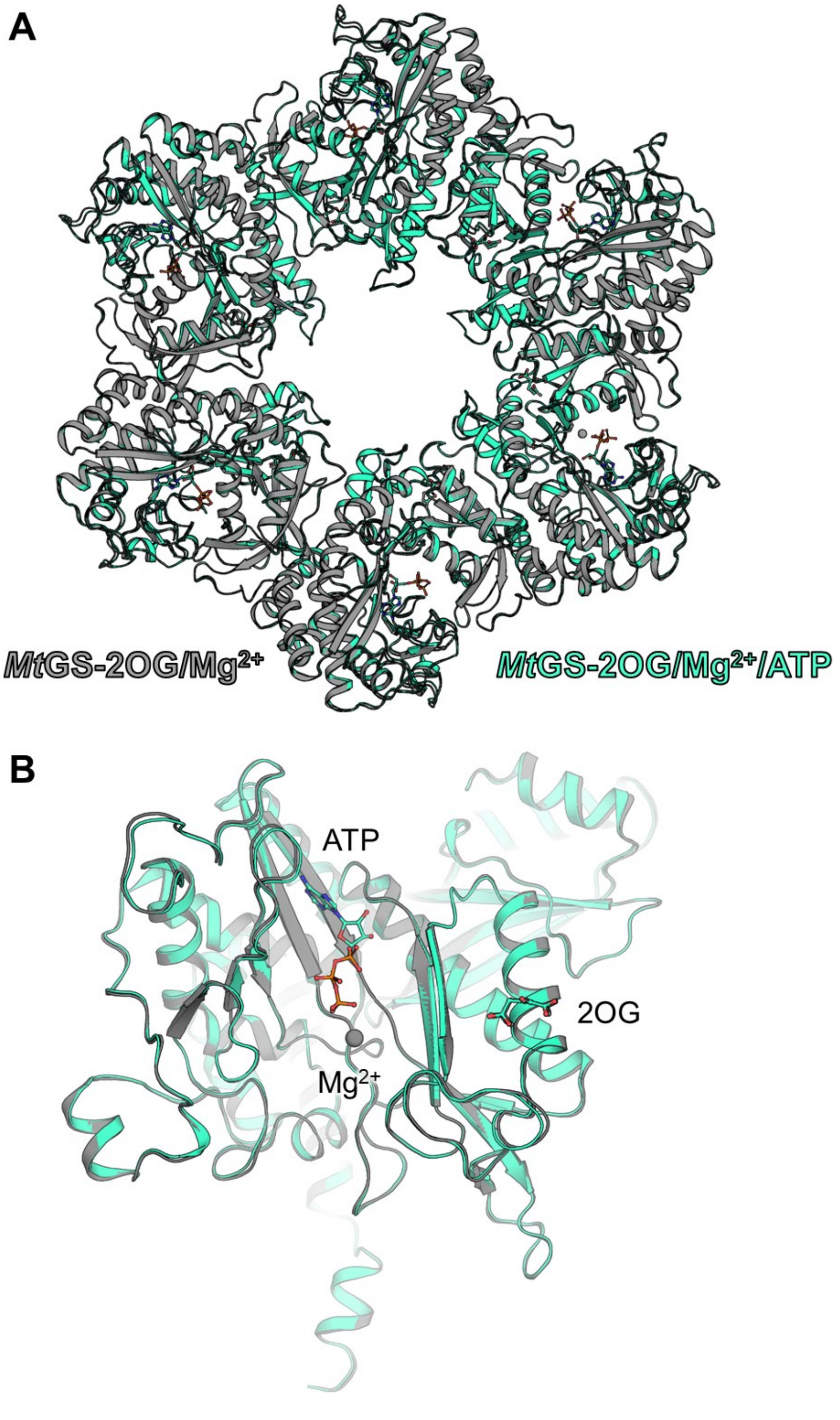


**Figure S8: Differences between 2OG/$Mg^{2+}$-containing and the 2OG/$Mg^{2+}$/ATP-containing *Mt*GS.** (A) Top view of the hexamer aligned on the C-terminal domain. (B) Side view of the monomer with the same pose as in Fig. 2C. Models are represented as cartoon with $Mg^{2+}$, 2OG and ATP as balls and sticks. Carbon, oxygen, nitrogen and phosphorous are colored in cyan, red, blue, and orange, respectively.

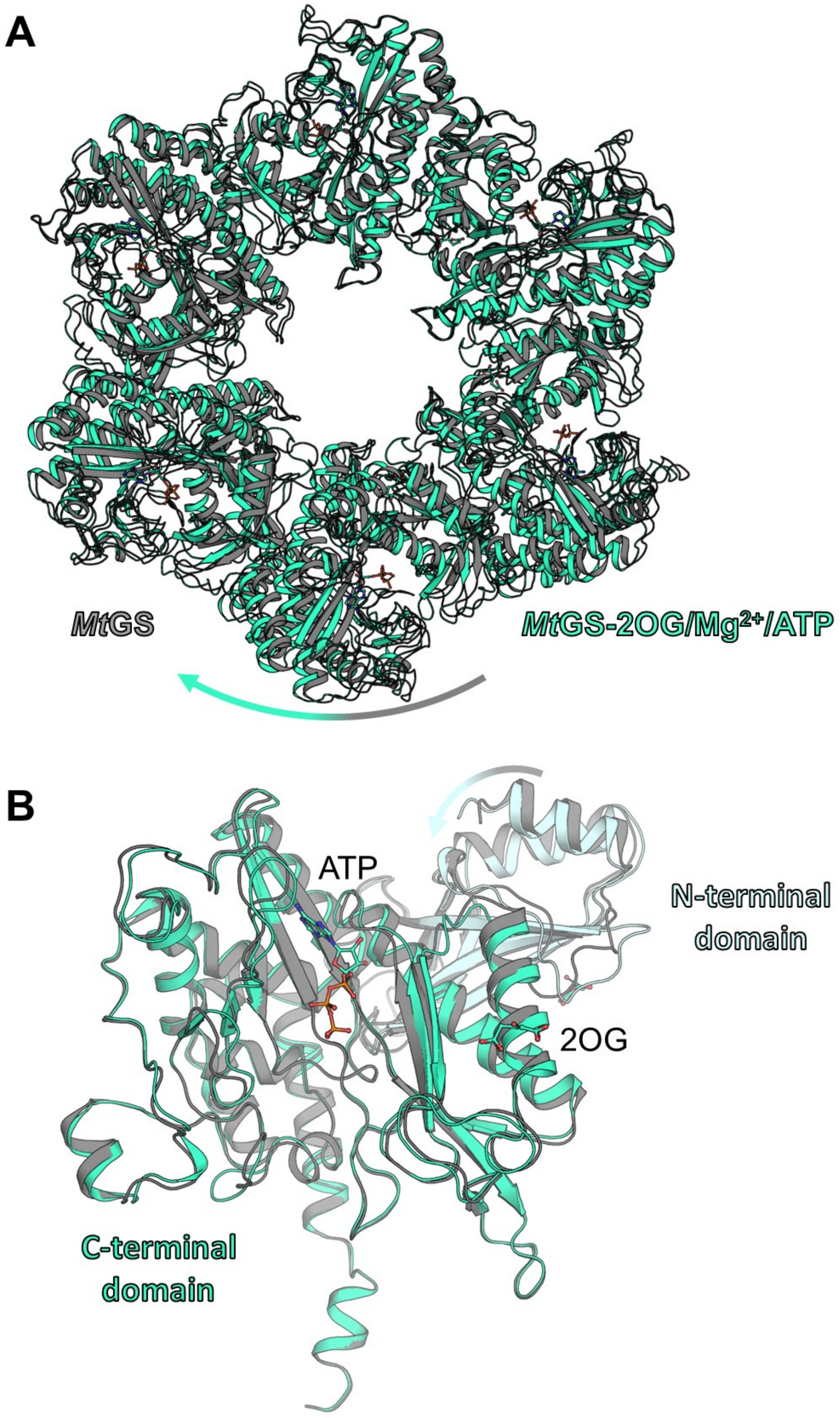


**Figure S9: Differences between apo and the 2OG/$Mg^{2+}$/ATP-containing *Mt*GS.** (A) Top view of the superposed *Mt*GS apo and *Mt*GS with 2OG/$Mg^{2+}$/ATP hexamers. (B) Side view of the monomer, aligned on the C-terminal domain with the same pose as in Fig. 2C. Models are represented as cartoons with 2OG, $Mg^{2+}$, and ATP as balls and sticks. Carbon, oxygen, nitrogen and phosphorous are colored in cyan, red, blue, and orange, respectively. Structural rearrangements are indicated with arrows.

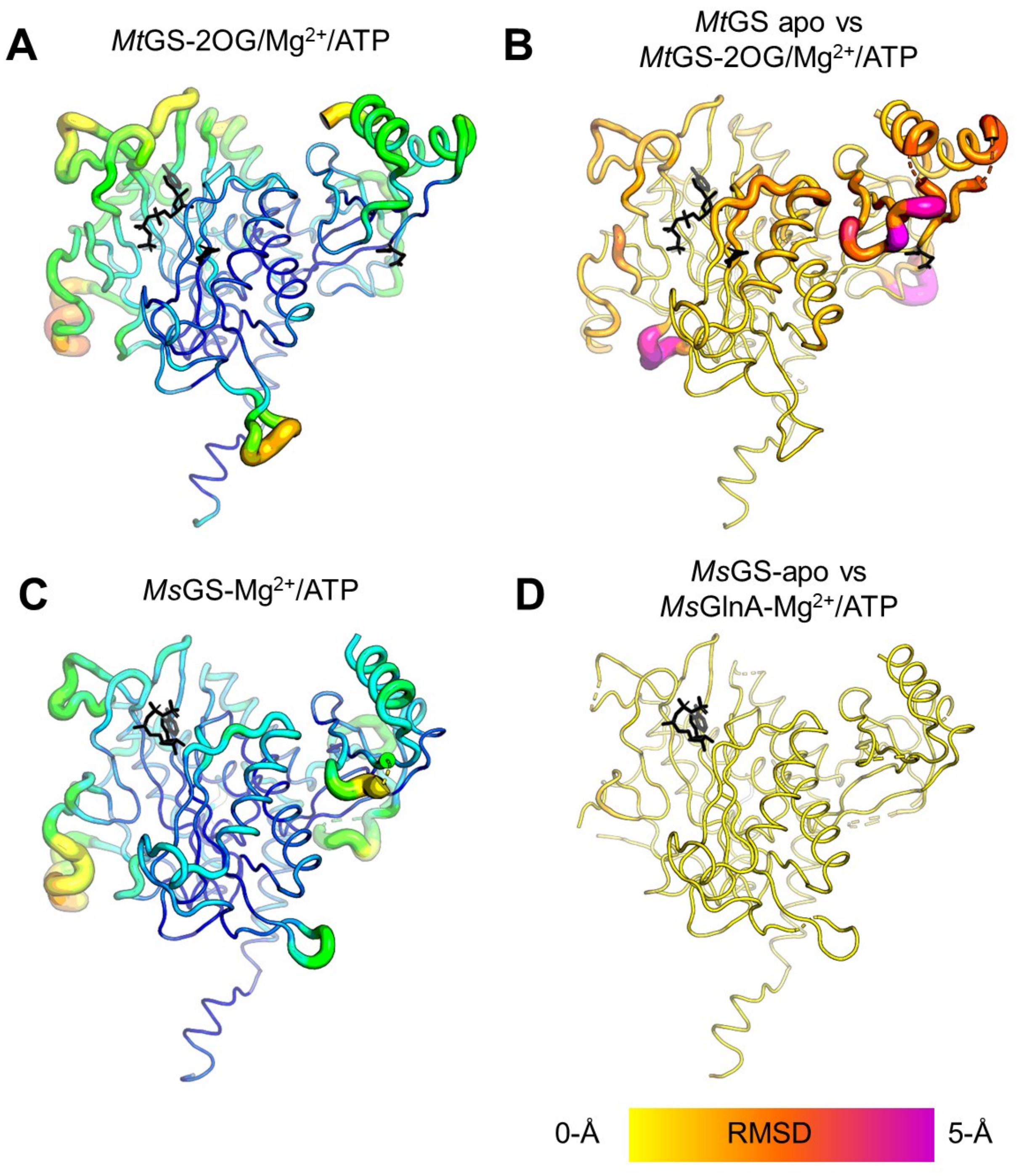


**Figure S10: Structural variation between *Mt*GS/*Ms*GS apo and ligand-bound state**. (A-B) B-factors and (C-D) RMSD values are displayed as putty. For the latter, the average of all chains is displayed. *Ms*GS $Mg^{2+}$/ATP and *Mt*GS 2OG/$Mg^{2+}$/ATP *versus* their respective apo state. Ligands are shown as black sticks.

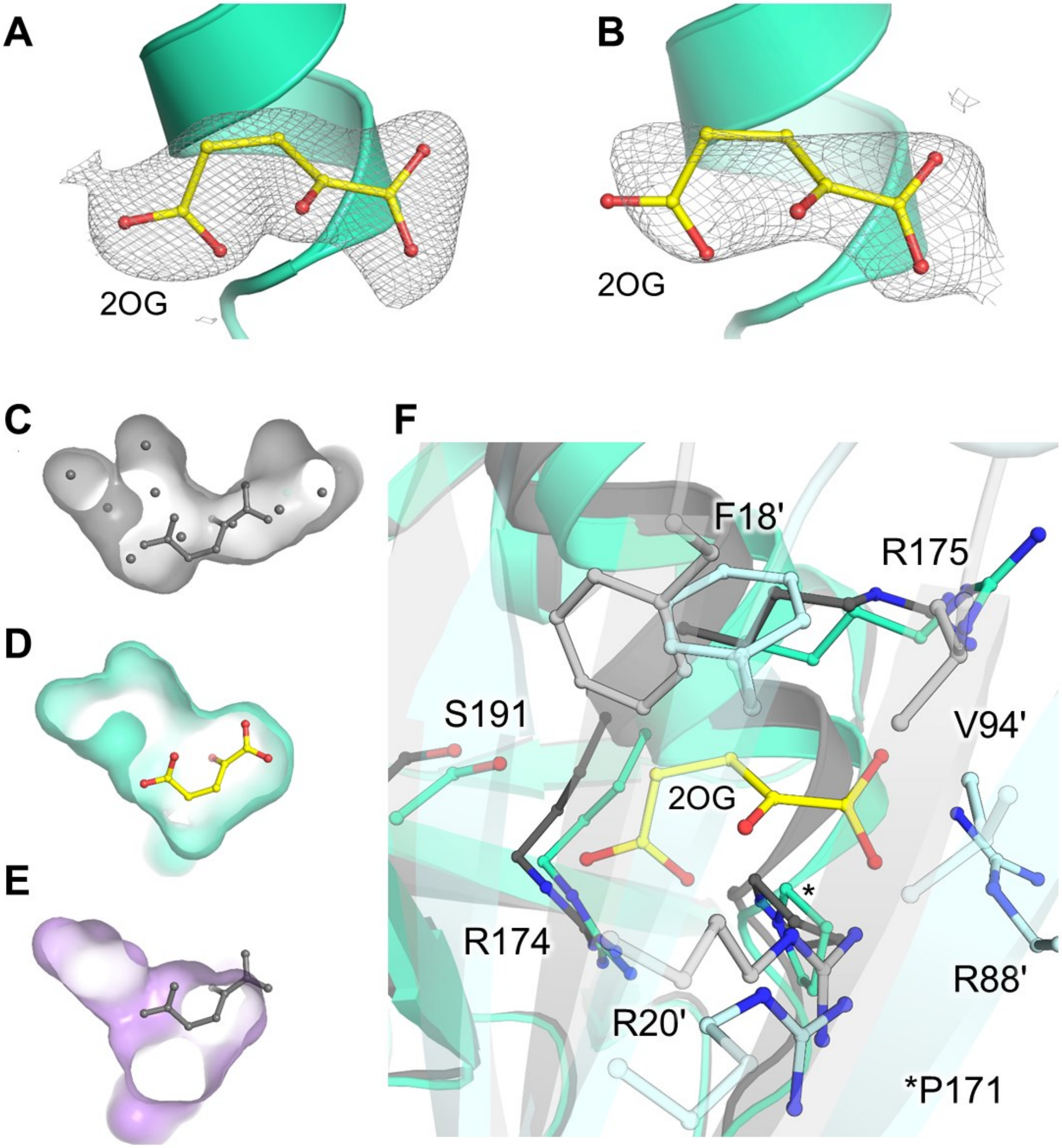


**Figure S11: 2OG binding site in *Mt*GS and *Ms*GS.** (A and B), Omit maps (2$F_o$-$F_c$ map contoured at 1 sigma, as grey mesh) of 2OG for *Mt*GS-2OG/$Mg^{2+}$/ATP (A) and *Mt*GS-2OG/$Mg^{2+}$ (B). 2OG as balls and sticks. (C-E) 2OG pocket surface in *Mt*GS apo (grey), with waters as grey spheres (C), *Mt*GS-2OG/$Mg^{2+}$/ATP (cyan) (D), and the equivalent position in *Ms*GS-$Mg^{2+}$/ATP (purple) (E). 2OG was superposed on the C-terminal domain from *Mt*GS-2OG/$Mg^{2+}$/ATP (grey balls and sticks) for panels C and E. (F) Overlay of *Mt*GS apo (grey) and *Mt*GS-2OG/$Mg^{2+}$/ATP (cyan). Residues in the close vicinity of 2OG and 2OG itself are shown as balls and sticks. The adjacent subunit is colored in a lighter shade. A-F are represented in cartoon. Oxygen and nitrogen are colored red and blue, respectively. Carbons are colored by chain and 2OG carbons in yellow.

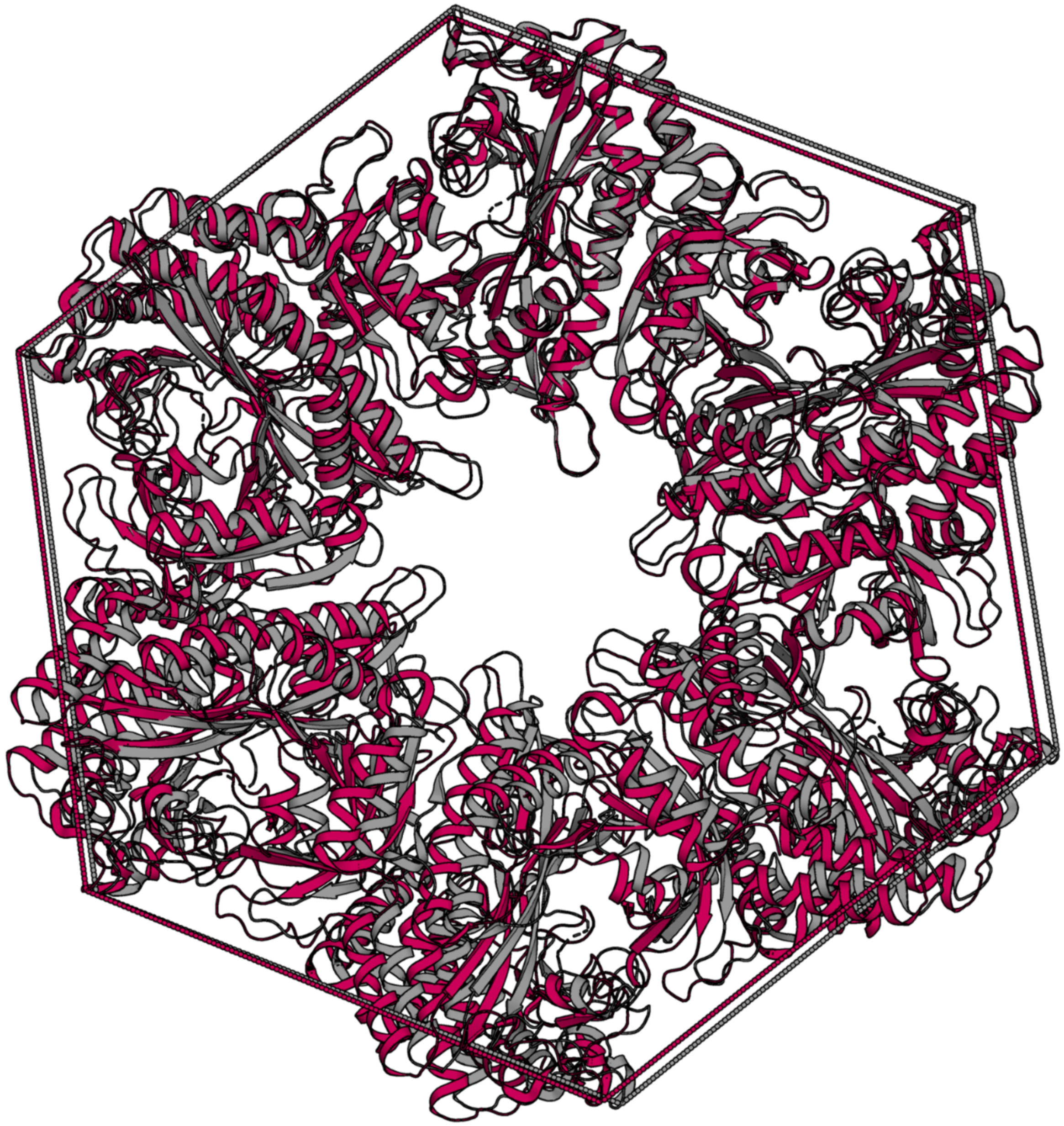

**Figure S12: Conformational change in *Bs*GS.** Superposition of the *Bs*GS apo (grey, PDB 4LNN) and *Bs*GS containing L-methionine-S-sulfoximine phosphate (SOX, which mimics the reaction intermediate, colored red, PDB 4LNI). Hexamers are represented as cartoons. The superposition was done on one monomer, and a dashed line was drawn on the Cα position of Asn262 to illustrate the overall movements.

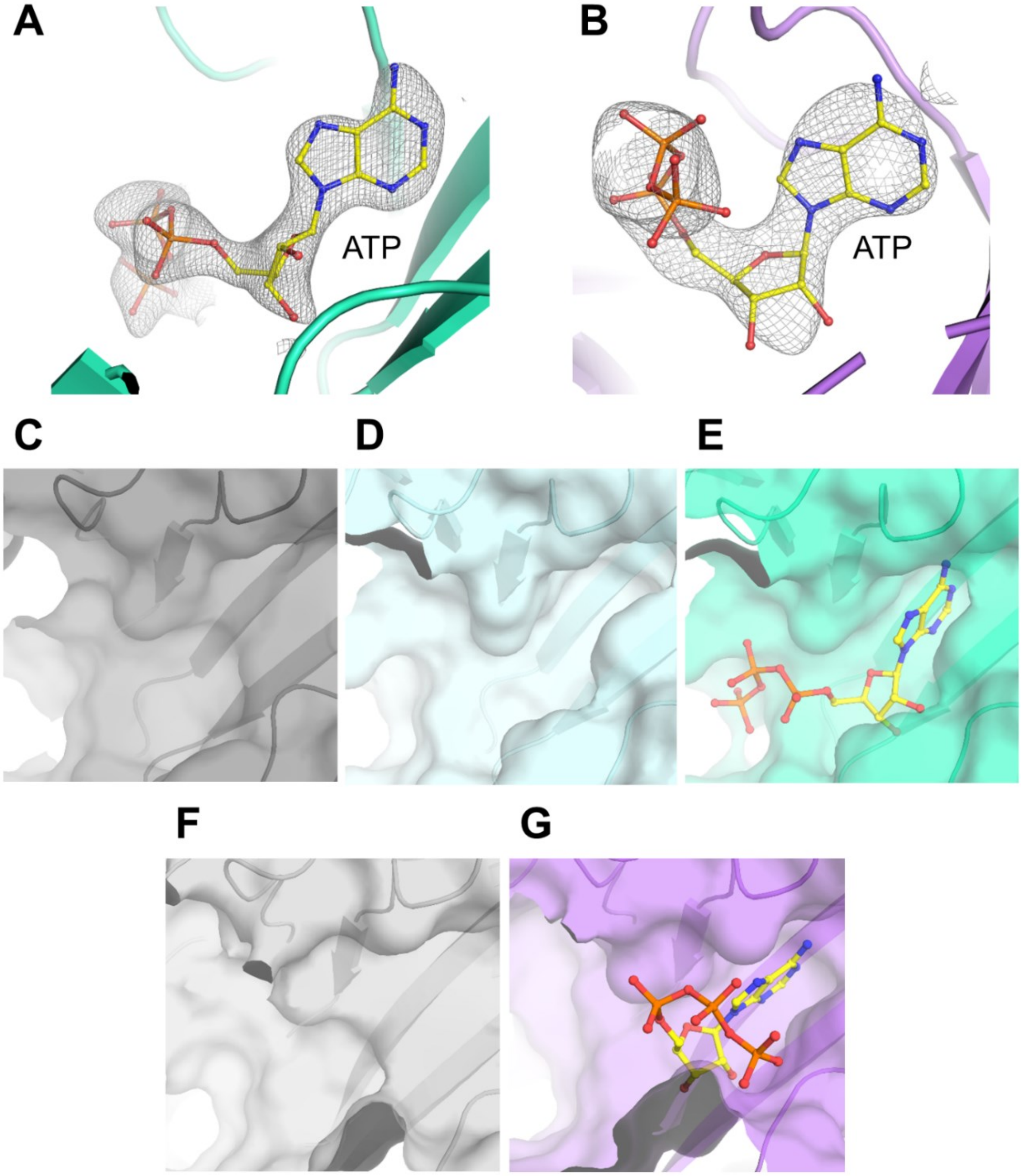


**Figure S13: ATP binding pocket in *Mt*GS and *Ms*GS.** (A-B) Omit maps (2$F_o$-$F_c$ contoured at 1 sigma, colored in grey mesh) of ATP for *Mt*GS-2OG/$Mg^{2+}$/ATP (A) and *Ms*GS-$Mg^{2+}$/ATP (B). (C-E) ATP pockets are displayed as transparent surfaces in *Mt*GS apo (C), *Mt*GS-2OG/$Mg^{2+}$ (D), and *Mt*GS-2OG/$Mg^{2+}$/ATP (E). (F and G) ATP pocket in *Ms*GS apo (F) and *Ms*GS-$Mg^{2+}$/ATP (G). ATP is highlighted as balls and sticks. Carbon, oxygen, nitrogen, and phosphorus are colored yellow, red, blue, and orange, respectively. All models are represented as cartoons.

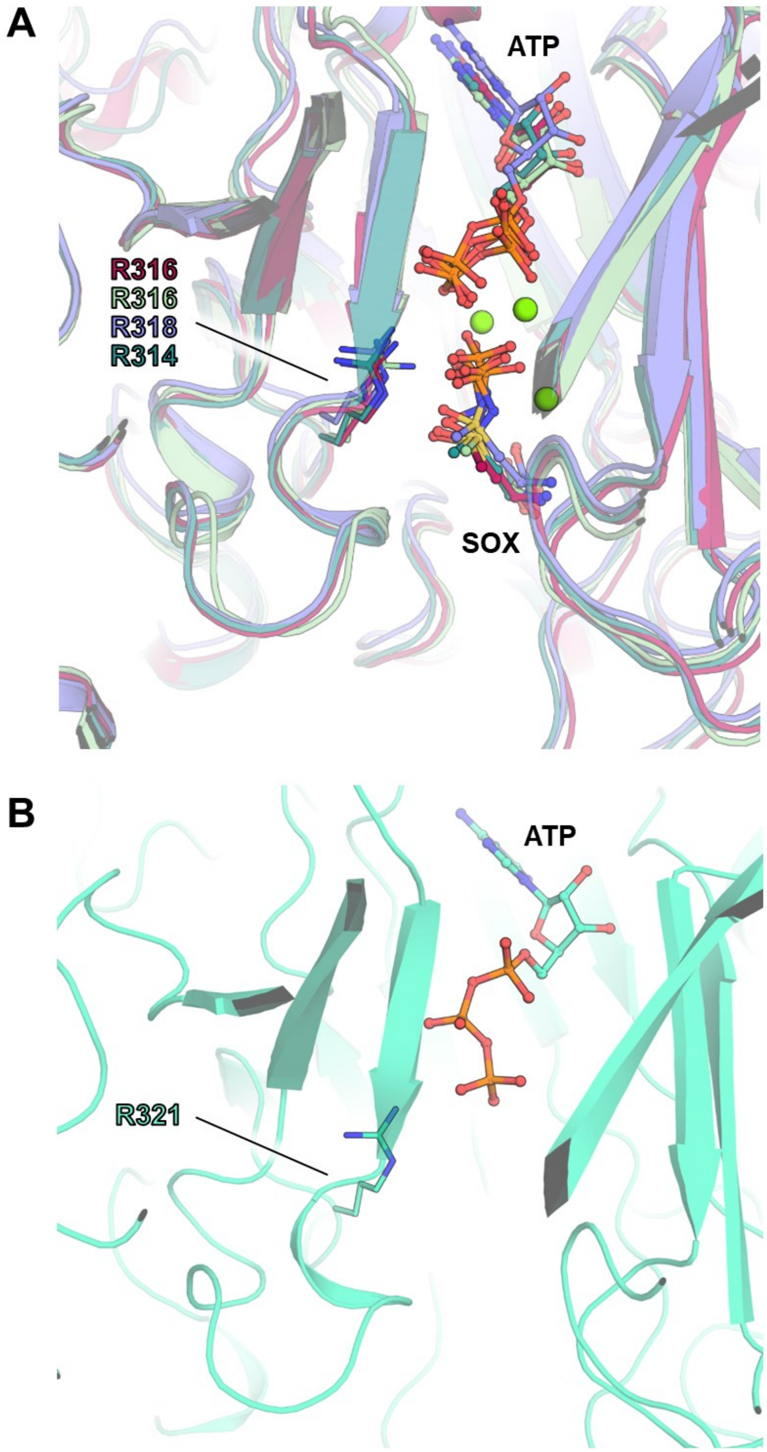


**Figure S14: Glutamate binding site view in all structurally characterized bacterial GSI-α and *Mt*GS.** (A) *Bs*GS (red, 4LNI), *Staphylococcus aureus* (blue, 7TDV), *Paenibacillus polymyxa* (dark green, 7TDP), *Listeria monocytogenes* (light green, 7TEN), all in the transition state with SOX/$Mg^{2+}$/ADP bound. (B) *Mt*GS. All models are in cartoons with ligands in balls and sticks. Oxygen, nitrogen, sulfur, phosphorus and magnesium are colored red, blue, yellow, orange and green, respectively. Carbons are accordingly colored to the model. The catalytic arginine is represented as sticks.

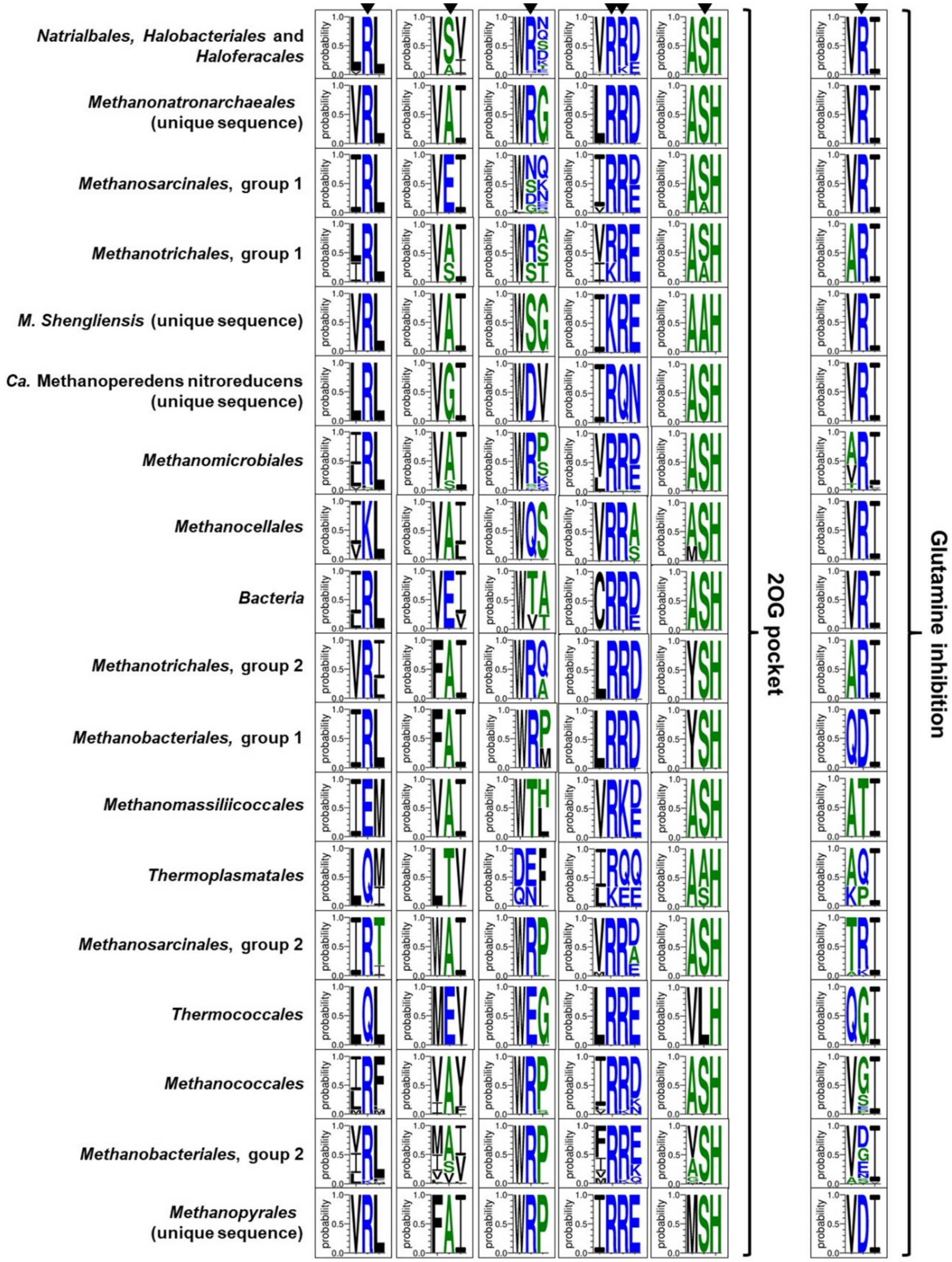


**Figure S15: Conservation of residues binding 2OG and glutamine in GSI-α.** The sequence conservation at the residues allowing or preventing binding of 2OG and glutamine (according to *Mt*GS, *Ms*GS and *Bs*GS and shown by black triangles) are illustrated by the Weblogo3 figures constructed on the sequences forming the GS monophyletic groups selected on the tree from Fig. 6.

1. Travis, B. A.; Peck, J. V.; Salinas, R.; Dopkins, B.; Lent, N.; Nguyen, V. D.; Borgnia, M. J.; Brennan, R. G.; Schumacher, M. A., Molecular dissection of the glutamine synthetase-GlnR nitrogen regulatory circuitry in Gram-positive bacteria. *Nat Commun* **2022,** *13* (1), 3793.
